# Rapid lethality of mice lacking the phagocyte oxidase and Caspase1/11 following *Mycobacterium tuberculosis* infection

**DOI:** 10.1101/2023.02.08.527787

**Authors:** Sean M. Thomas, Andrew J. Olive

**Affiliations:** Department of Microbiology and Molecular Genetics, College of Osteopathic Medicine, Michigan State University, East Lansing, MI USA

## Abstract

Immune networks that control antimicrobial and inflammatory mechanisms have overlapping regulation and functions to ensure effective host responses. Genetic interaction studies of immune pathways that compare host responses in single and combined knockout backgrounds are a useful tool to identify new mechanisms of immune control during infection. For disease caused by pulmonary *Mycobacterium tuberculosis* infections, which currently lacks an effective vaccine, understanding genetic interactions between protective immune pathways may identify new therapeutic targets or disease-associated genes. Previous studies suggested a direct link between the activation of NLRP3-Caspase1 inflammasome and the NADPH-dependent phagocyte oxidase complex during Mtb infection. Loss of the phagocyte oxidase complex alone resulted in increased activation of Caspase1 and IL1β production during Mtb infection, resulting in failed disease tolerance during the chronic stages of disease. To better understand this interaction, we generated mice lacking both *Cybb*, a key subunit of the phagocyte oxidase, and *Caspase1/11*. We found that *ex vivo* Mtb infection of *Cybb*^*-/-*^*Caspase1/11*^*-/-*^ macrophages resulted in the expected loss of IL1β secretion but an unexpected change in other inflammatory cytokines and bacterial control. Mtb infected *Cybb*^*-/-*^*Caspase1/11*^*-/-*^ mice rapidly progressed to severe TB, succumbing within four weeks to disease characterized by high bacterial burden, increased inflammatory cytokines, and the recruitment of granulocytes that associated with Mtb in the lungs. These results uncover a key genetic interaction between the phagocyte oxidase complex and Caspase1/11 that controls protection against TB and highlight the need for a better understanding of the regulation of fundamental immune networks during Mtb infection.

## Introduction

Defense against infection requires the regulated activation of immune networks that determine the magnitude and duration of the host response (1, 2). Dysregulation of these immune networks contributes to increased susceptibility to infection and reduced disease tolerance (3-5). During lung infections with *Mycobacterium tuberculosis*, pro-inflammatory responses mediated by cytokines such as interleukin 1−beta (IL1β), tumor necrosis factor (TNF) and interferon-gamma (IFNγ) must be strong enough to restrict infection, while maintaining respiratory function and controlling tissue damage (5-9). This balance is controlled by the tight regulation of cytokine and chemokine secretion to effectively direct the inflammatory process and immune cell recruitment (10, 11). Disruption of this balance contributes to progressive inflammatory tuberculosis (TB) disease which results in over 1.5 million deaths each year (12). Understanding the factors contributing to protection or susceptibility during Mtb infection is essential to devise more effective therapies and immunization strategies.

TB susceptibility is controlled by a combination of bacterial, host and environmental factors (13, 14). Many defined protective host genes comprise the mendelian susceptibility to mycobacterial diseases (MSMD) (15). Patients with these conditions have loss-of-function alleles in genes that are essential for protective host responses such as IFNγ signaling. Additional genes related to autophagy, reactive oxygen and nitrogen species (ROS/RNS) production, and cytokine production are also protective in the mouse model of Mtb (16-19). However, while many genes are now identified as protective against Mtb, the precise mechanisms by which they control disease remains unclear.

One such protective mechanism is the ROS produced by the NADPH Phagocyte Oxidase (20). In humans, Chronic Granulomatous Disease (CGD) in patients with dysfunctional phagocyte oxidase complexes is associated with increased susceptibility to mycobacterial infections (21). Mice deficient in the phagocyte oxidase subunit *Cybb* control Mtb replication yet show defects in disease tolerance that result in a modest reduction in survival following high dose Mtb infection (18, 22-24). The loss of *Cybb* results in the hyperactivation of the NLRP3 inflammasome and exacerbated IL1β production by bone marrow-derived macrophages (BMDMs) and *in vivo* during murine Mtb infection (18). This exacerbated IL1β can be reversed in BMDMs with chemical inhibitors of NLRP3 or Caspase1. Caspase1 is a critical component of the NLRP3 inflammasome and is responsible for the activation of mature IL1β, IL18, and Gasdermin D (25, 26). However, while Mtb infection of Caspase1-deficient macrophages results in loss of mature IL1β production, mice lacking Caspase1/11 have no defects in IL1β and minimal changes in susceptibility to TB *in vivo* (27, 28). Even though previous studies found clear links between phagocyte oxidase and the NLRP3 inflammasome that contribute to protection during Mtb infection, how these pathways interact and regulate each other’s function remains unclear.

Here, we used a genetic approach to understand interactions between the phagocyte oxidase and the inflammasome by generating *Cybb*^*-/-*^*Caspase1/11*^*-/-*^ animals. Mtb infection of macrophages and dendritic cells from these animals reversed the exacerbated IL1β production that was responsible for failed tolerance in *Cybb*^-/-^ cells. However, we found dysregulation of other pro-inflammatory mediators and reduced bacterial control during infection of *Cybb*^*-/-*^*Caspase1/11*^*-/-*^ BMDMs. *In vivo*, we uncovered a synthetic susceptibility with *Cybb*^*-/-*^*Caspase1/11*^*-/-*^ animals succumbing rapidly to TB disease within 4 weeks. We observed the loss of bacterial control and the recruitment of permissive granulocytes in *Cybb*^*-/-*^ *Caspase1/11*^*-/-*^ that were not seen in wild type, *Cybb*^*-/-*^ or *Caspase1/11*^*-/-*^ animals. Thus, our results uncovered a previously unknown genetic interaction between the phagocyte oxidase and the Caspase1 inflammasome that contributes to TB protection. Furthermore, our results highlight the complexity of the interactions between immune networks that control Mtb susceptibility and the importance of the regulation of inflammatory cytokines in the lung environment.

## Results

### Loss of Caspase 1/11 results in decreased IL1β production in *Cybb*^*-/-*^ phagocytes

Macrophages deficient in the phagocyte oxidase subunit *Cybb* hyperactivate the NLRP3 inflammasome and produce damaging levels of IL1β during Mtb infection (18). We developed a genetic model to understand the interaction between these genes by generating mice deficient in both *Cybb* and *Caspase1/11* in the C57BL6/J background. We first examined the regulation of IL1β during Mtb infection in cells lacking *Cybb*^*-/-*^*Caspase1/11*^*-/-*^. Bone marrow-derived macrophages (BMDMs) from wild type, *Cybb*^*-/-*^, *Caspase1/11*^*-/-*^, and *Cybb*^*-/-*^*Caspase1/11*^*-/-*^ mice were infected with Mtb H37Rv. 14 hours later, the supernatants were removed from infected and uninfected control cells and the levels of IL1β were quantified by ELISA. As previously shown, Mtb infected *Cybb*^*-/-*^ phagocytes secreted significantly more IL1β compared to wild type cells while *Caspase1/11*^*-/-*^ cells released nearly undetectable levels of IL1β (Figure 1A) (18). Loss of Caspase1/11 in combination with Cybb resulted in no IL1β release, similar to what was observed in *Caspase1/11*^*-/-*^ macrophages. The experiment was repeated using bone marrow-derived dendritic cells (BMDCs) and the results were consistent with BMDMs. *Cybb*^*-/-*^ cells produce high levels of IL1β which is reversed in the absence of Caspase1/11 (Figure 1B). These data show that loss of Caspase1/11 reverses the elevated IL1β production observed in *Cybb*^*-/-*^ deficient phagocytes infected with Mtb.

**Figure 1.**
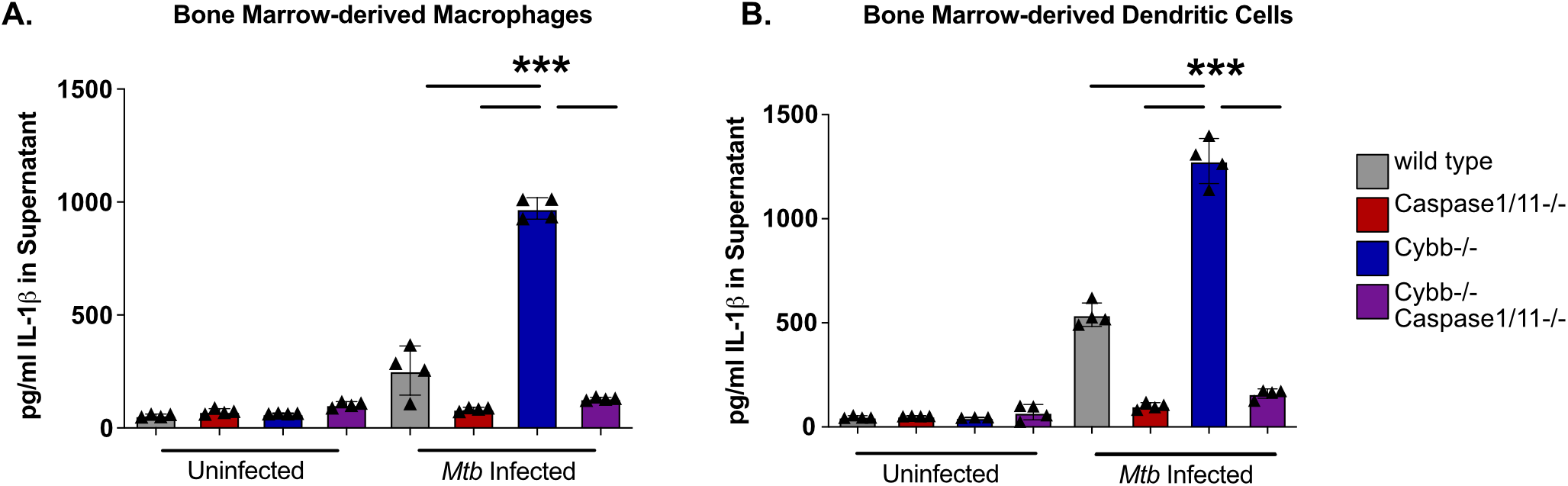
Exacerbated IL1β following Mtb infection of *Cybb*^*-/-*^ myeloid-cells is dependent on Caspase1/11. **(A)** BMDMs or **(B)** BMDCs from wild type, *Caspase1/11*^*-/-*^, *Cybb*^*-/-*^ and *Cybb*^*-/-*^ *Caspase1/11*^*-/-*^ mice were left uninfected or infected with Mtb H37Rv at an MOI of 5. The following day IL1β was quantified from the supernatants by ELISA. Each point represents data from a single well from one representative experiment of three. *** p<.001 by one-way ANOVA with a tukey test for multiple comparisons.

### Cybb^-/-^Caspase1/11^-/-^ BMDMs dysregulate cytokines and Mtb control during infection

Both the ROS produced by the phagocyte oxidase and the immune pathways regulated by the inflammasome can modulate the inflammatory state of macrophages (29, 30). To better understand how the functions of *Cybb* and *Caspase1/11* interact to regulate inflammation, we infected BMDMs from each genotype with Mtb and we quantified cell death and cytokine release via multiplex cytokine analysis. Over the 14-hour infection, we observed no significant differences in cell death between any genotype (Figure 2A). Similar to the ELISA above, we observed increased IL1β production by *Cybb*^*-/-*^ macrophages which was reversed in macrophages from *Cybb*^*-/-*^*Caspase1/11*^*-/-*^ mice (Figure 2B). While IL1α production was also increased by Cybb^-/-^ cells, this was not reversed and was, in contrast to IL1β, exacerbated in *Cybb*^*-/-*^*Caspase1/11*^*-/-*^ macrophages. The increased IL1α production was not due to loss of Caspase1/11 alone, since *Caspase1/11*^*-/-*^ BMDMs produced nearly undetectable levels of IL1α following Mtb infection. Thus, IL1α production by BMDMs is exacerbated in the absence of both *Cybb* and *Caspase1/11*.

**Figure 2.**
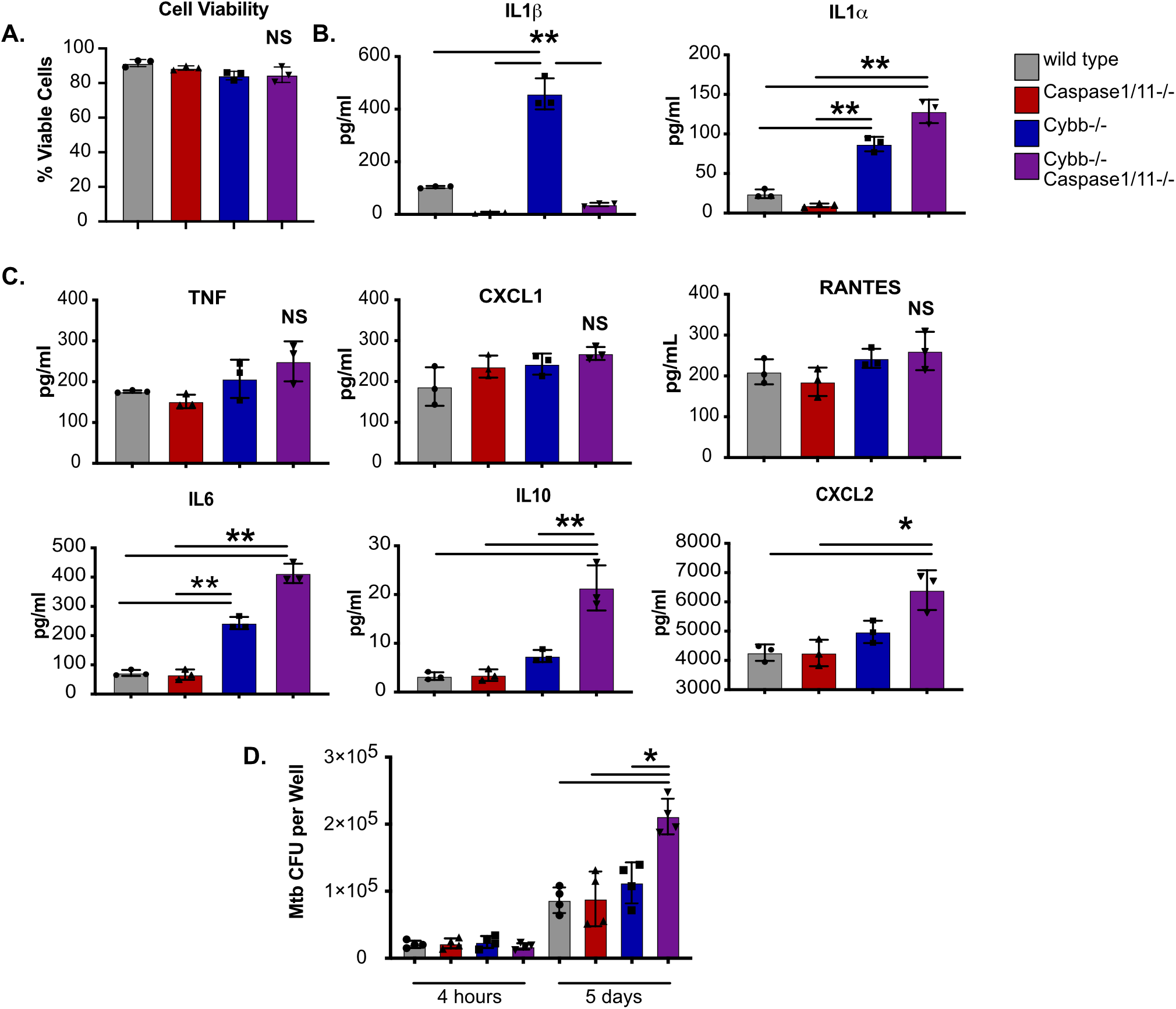
*Cybb*^*-/-*^*Caspase1/11*^*-/-*^ macrophages are hyperinflammatory and permissive to bacterial growth during Mtb infection. **(A)** BMDMs from wild type, *Caspase1/11*^*-/-*^, *Cybb*^*-/-*^ and *Cybb*^*-/-*^ *Caspase1/11*^*-/-*^ mice were left uninfected or were infected with Mtb H37Rv at an MOI of 5. The following day, total viable cells were quantified in each infection condition and normalized to uninfected control cells. Shown is percent viability of infected cells compared to uninfected cells of the same genotype. **(B)** BMDMs from wild type, *Caspase1/11*^*-/-*^, *Cybb*^*-/-*^ and *Cybb*^*-/-*^*Caspase1/11*^*-/-*^ mice were infected with Mtb H37Rv at an MOI of 5. The following day, cytokines from the supernatant were quantified by Luminex multiplex assay. Shown are results for IL1β and IL1α, and **(C)** other indicated cytokines (TNF, CXCL1, RANTES, IL6, IL10, and CXCL2). **(D)** BMDMs from wild type, *Caspase1/11*^*-/-*^, *Cybb*^*-/-*^ and *Cybb*^*-/-*^ *Caspase1/11*^*-/-*^ mice were infected with Mtb H37Rv at an MOI of 1. At the indicated timepoints, cells were lysed and viable Mtb CFU were quantified. In all experiments, each point represents data from a single well and shown is mean ^+^/- SD from one representative experiment of two or three similar experiments. * p<.05 ** p<.01 NS no significance, by one-way ANOVA with a tukey test for multiple comparisons.

The multiplex cytokine panel included a range of other inflammatory cytokines that were compared between each macrophage genotype (Figure 2C). Most cytokines, including TNF, RANTES and CXCL1 showed no significant difference between any of the genotypes. Most cytokines, including TNF, RANTES and CXCL1 showed no significant difference between any of the genotypes. However, IL6 and IL10 production were both significantly increased by *Cybb*^*-/-*^ BMDMs which was further exacerbated by *Cybb*^*-/-*^*Caspase1/11*^*-/-*^ cells. Finally, CXCL2 was significantly increased only in *Cybb*^*-/-*^ *Caspase1/11*^*-/-*^ macrophages. Taken together, *Cybb*^*-/-*^*Caspase1/11*^*-/-*^ macrophages dysregulate a range of inflammatory cytokines in response to Mtb infection.

Since the inflammatory milieu was altered during infection of *Cybb*^*-/-*^*Caspase1/11*^*-/-*^ BMDMs, we next tested if intracellular control of Mtb growth was compromised. BMDMs from each genotype were infected with Mtb and growth was monitored using a CFU assay. We observed no significant difference between genotypes in bacterial uptake 4 hours following infection (Figure 2D). 5 days later we observed no change in bacterial control in *Cybb*^*-/-*^ or *Caspase1/11*^*-/-*^ BMDMs but found significantly more Mtb growth in *Cybb*^*-/-*^*Caspase1/11*^*-/-*^ BMDMs. These data suggest that the loss of *Cybb* and *Caspase1/11* together does not compromise cell survival but does result in less effective Mtb control and dysregulated cytokine production that does not occur in either knockout mouse genotype alone.

### *Cybb*^*-/-*^*Caspase1/11*^*-/-*^ mice are hyper-susceptible to Mtb infection

Our experiments in BMDMs showed that the loss of *Cybb* and *Caspase1/11* together results in dysregulated host responses during Mtb infection. We hypothesized that this dysregulation would result in changes to *in vivo* TB disease progression. To test this hypothesis, wild type, *Cybb*^*-/-*^, *Caspase1/11*^*-/-*^, and *Cybb*^*-/-*^*Caspase1/11*^*-/-*^ mice were infected with Mtb by low dose aerosol. As mice were monitored during the infection, we observed dramatic weight loss of *Cybb*^*-/-*^*Caspase1/11*^*-/-*^ animals that required almost all animals to be euthanized prior to 30 days post-infection (Figure 3A). In contrast, all other genotypes had gained weight over the same time of infection. Survival analysis during these infections found that *Cybb*^*-/-*^*Caspase1/11*^*-/-*^ mice are highly susceptible to Mtb infection, with all animals requiring euthanasia earlier than 5 weeks post infection (Figure 3B). In contrast, wild type, *Cybb*^*-/-*^ and *Caspase1/11*^*-/-*^ animals all survived beyond day 75 similar to previous studies (18, 27, 28). This observation suggests a strong genetic interaction between *Cybb* and *Caspase1/11* that results in the synthetic hyper-susceptibility of animals to *Mtb* infection.

**Figure 3.**
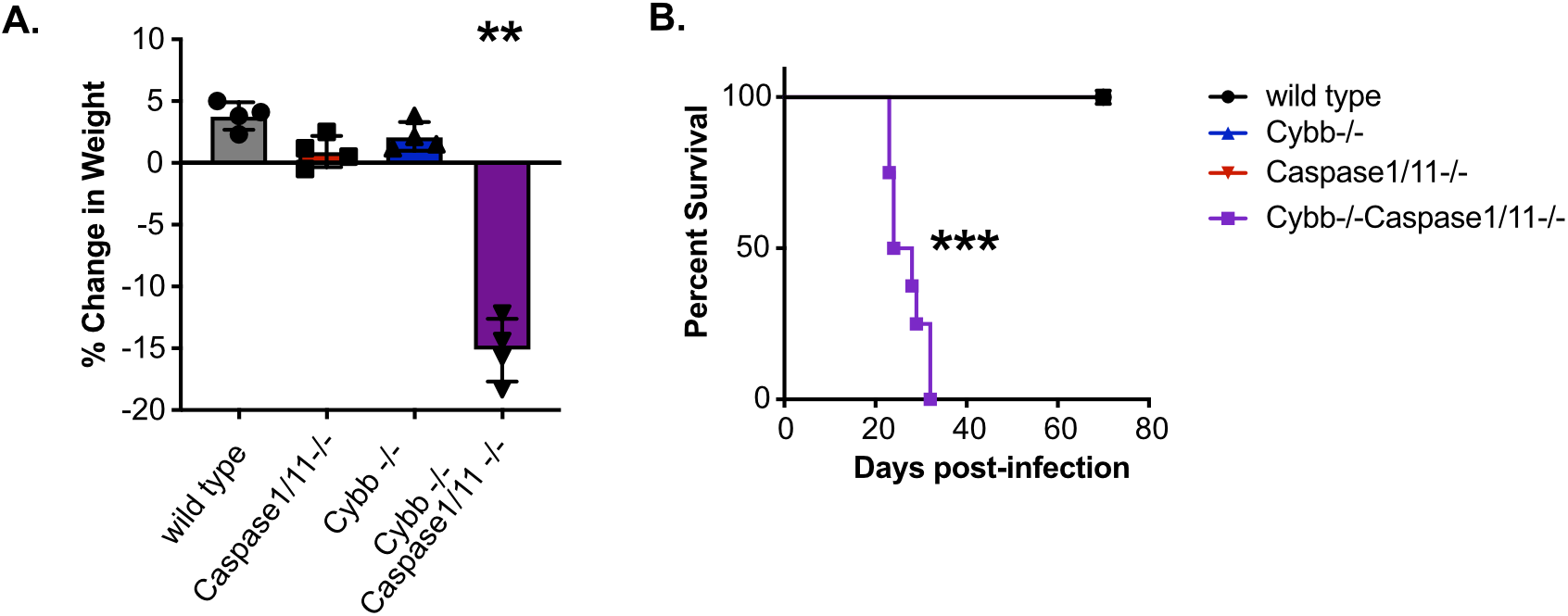
*Cybb*^*-/-*^*Caspase1/11*^*-/-*^ mice rapidly succumb to pulmonary Mtb infection. Wild type, *Caspase1/11*^*-/-*^, *Cybb*^*-/-*^ and *Cybb*^*-/-*^*Caspase1/11*^*-/-*^ mice were infected with Mtb H37Rv by the aerosol route in a single batch (Day 1 50-150 CFU). **(A)** Change in mouse weight from Day 0 to 24 days post-infection was quantified. Data are from one experiment and are representative of three similar experiments. Statistics were determined by a Mann Whitney test **p<.01. **(B)** The relative survival of each genotype was quantified over 75 days of infection. Data are pooled from two independent experiments. Statistics were determined by a Mantel-Cox test *** p<.001.

### Mtb infection of *Cybb*^*-/-*^*Caspase1/11*^*-/-*^ mice results in increased bacterial growth and inflammatory cytokine production

We next sought to determine the mechanisms driving the susceptibility of *Cybb*^*-/-*^*Caspase1/11*^*-/-*^ animals. Wild type, *Cybb*^*-/-*^, *Caspase1/11*^*-/-*^, and *Cybb*^*-/-*^*Caspase1/11*^*-/-*^ mice were infected with H37Rv YFP by low dose aerosol, and 25 days later, viable Mtb in the lungs and spleen were quantified by CFU plating (31). We observed similar numbers of Mtb in wild type, *Cybb*^*-/-*^, and *Caspase1/11*^*-/-*^ animals in both organs and in line with previous reports (18, 27, 28). In contrast, over 10-fold more Mtb were present in the lungs and ~5-fold more Mtb were present in the spleens of infected *Cybb*^*-/-*^*Caspase1/11*^*-/-*^ mice (Figure 4A and 4B). We further characterized the cytokine profile from infected lung homogenates using a Luminex multiplex assay. We found that *Cybb*^*-/-*^*Caspase1/11*^*-/-*^ mice express high levels of inflammatory cytokines including IL1α, IL1β, TNF, and IL6 but not IL10 (Figure 4C). We observed no significant differences between *Caspase1/11*^*-/-*^ and wild type mice, while in *Cybb*^*-/-*^ mice we found increased levels of IL1β but no other cytokines in line with previous studies (18, 27, 28). Thus, mice deficient in both *Cybb* and *Caspase 1/11* are unable to effectively control Mtb replication and display hyperinflammatory cytokine responses.

**Figure 4.**
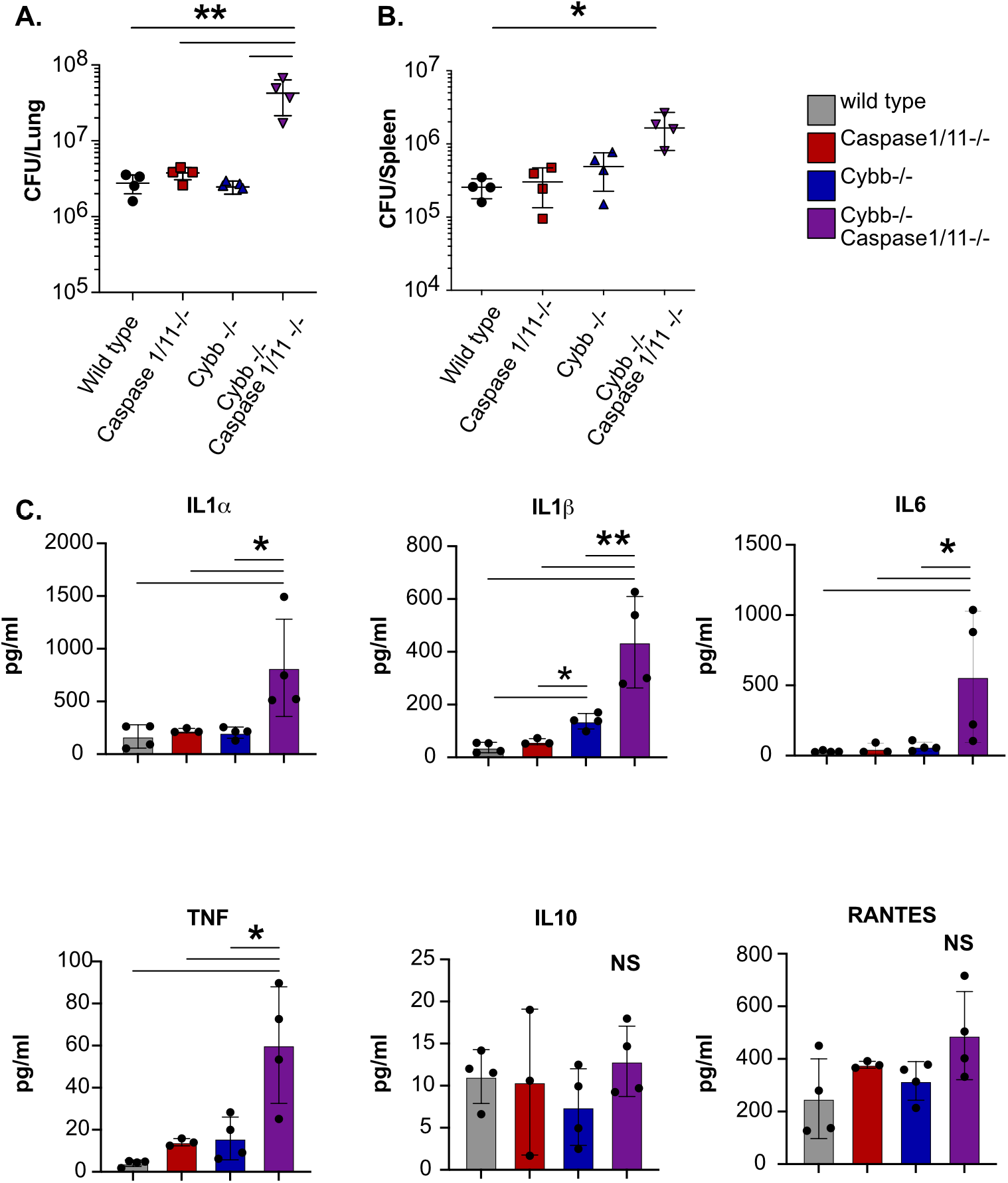
*Cybb*^*-/-*^*Caspase1/11*^*-/-*^ mice do not control Mtb growth and are hyperinflammatory. Wild type, *Caspase1/11*^*-/-*^, *Cybb*^*-/-*^ and *Cybb*^*-/-*^*Caspase1/11*^*-/-*^ mice were infected with Mtb H37Rv YFP by the aerosol route in a single batch (Day 1 50-100 CFU). Lungs and spleen were collected at 25 days post-infection and used to quantify bacterial CFU. **(A)** Bacterial burden in the lungs and **(B)** spleens of mice are shown. **(C)** Concentrations of cytokines in lung homogenates from infected mice were quantified (IL1α, IL1β, IL6, TNF, IL10 and RANTES). Each point represents a single mouse, data are representative of one experiment from three similar experiments. * p<.05 ** p<.01 NS no significance, by one-way ANOVA with a tukey test for multiple comparisons.

### Permissive granulocytes are recruited to the lungs of *Cybb*^*-/-*^*Caspase1/11*^*-/-*^ mice during Mtb infection

The extreme susceptibility and increased Mtb growth observed in Cybb^-/-^Caspase1/11^-/-^ mice is similar to mice lacking *IFNγ* or *Nos2 (7, 31-33)*. Recent work showed that the susceptibility of *Nos2*^-/-^ animals is driven by dysregulated inflammation that recruits permissive granulocytes to the lungs which then allow for amplified Mtb replication (33, 34). We hypothesized that similar responses may be associated with the susceptibility of *Cybb*^*-/-*^*Caspase1/11*^*-/-*^ mice during Mtb infection. To test this hypothesis, we first analyzed the myeloid-derived populations of cells in the lungs and spleen of wild type, *Caspase1/11*^*-/-*^ *Cybb*^*-/-*^ and *Cybb*^*-/-*^*Caspase1/11*^*-/-*^ animals infected with Mtb H37Rv YFP by low dose aerosol for 25 days. While wild type and *Caspase1/11*^*-/-*^ animals showed indistinguishable distributions of cells, *Cybb*^*-/-*^ mice recruited more GR1^hi^ CD11b^+^ neutrophils in agreement with our previous findings (Figure 5A and 5B) (18). However, we observed a significant increase in the total number of GR1^int^ CD11b^+^ cells in the lungs of *Cybb*^*-/-*^*Caspase1/11*^*-/-*^ mice. This population is consistent with the permissive myeloid cells seen in mice that are highly susceptible to Mtb infection (33, 34).

**Figure 5.**
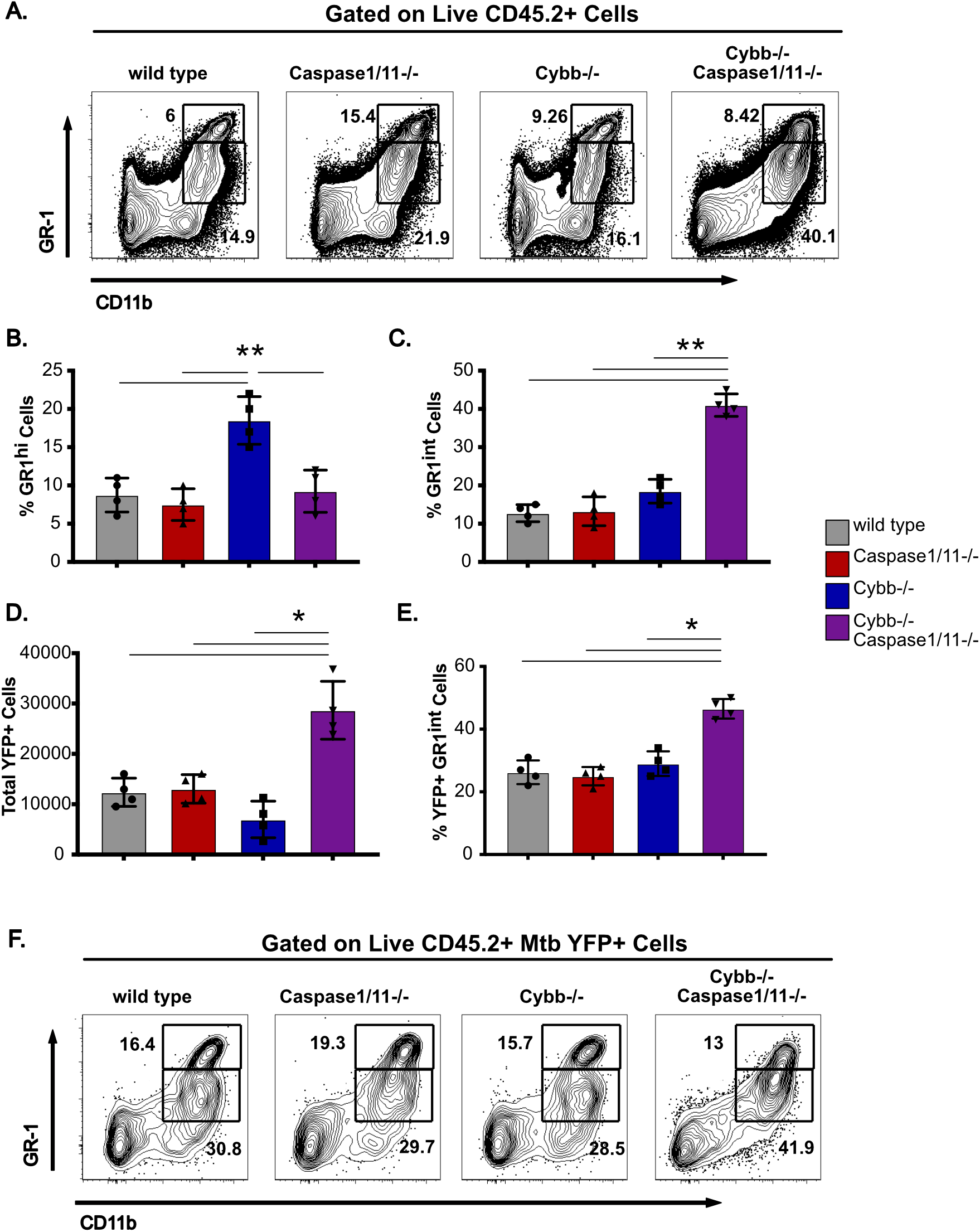
GR-1^int^ granulocytes are recruited to the lungs and are associated with Mtb during infection of *Cybb*^*-/-*^*Caspase1/11*^*-/-*^ mice. Wild type, *Caspase1/11*^*-/-*^, *Cybb*^*-/-*^ and *Cybb*^*-/-*^*Caspase1/11*^*-/-*^ mice were infected with Mtb H37Rv YFP by the aerosol route in a single batch (Day 1 50-100 CFU). Lungs were collected at 25 days post-infection and single cell homogenates were made for flow cytometry analysis. **(A)** Shown is a representative flow cytometry plot of total lung granulocytes based on CD11b and GR1 staining (Gated on live CD45.2^+^ single cells). Gates indicate CD11b^+^ GR1^hi^ or CD11b^+^ GR1^int^ granulocytes present in the lungs. **(B)** The percent of gated cells (live CD45.2^+^ single cells) that were CD11b^+^ GR1^hi^ and **(C)** CD11b^+^ GR1^int^ were quantified. **(D)** The total number of H37Rv YFP^+^ cells were quantified from each mouse lung following gating on live, CD45.2^+^ single cells. **(E)** The percent of gated H37Rv YFP^+^ cells that were CD11b^+^ GR1^int^ were quantified. **(F)** Shown is a representative flow cytometry plot of H37Rv YFP^+^ infected granulocytes (Gated on live CD45.2^+^ YFP^+^ single cells). Gates indicate CD11b^+^ GR1^hi^ or CD11b^+^ GR1^int^ granulocytes present in the lungs. Each point represents a single mouse and data are representative of one experiment from three similar experiments. * p<.05 ** p<.01 by one-way ANOVA with a tukey test for multiple comparisons.

If the recruited GR1^int^ CD11b^+^ granulocytes in the lungs of *Cybb*^*-/-*^*Caspase1/11*^*-/-*^ mice are permissive for Mtb growth, we predicted these cells would harbor a disproportionate fraction of intracellular Mtb in the lungs. To test this prediction, we quantified the total YFP^+^ infected cells from each genotype. We found an increase in the total number YFP^+^ cells only in *Cybb*^*-/-*^*Caspase1/11*^*-/-*^ mice (Figure5D). These data show that the lungs of *Cybb*^*-/-*^*Caspase1/11*^*-/-*^ mice harbor more infected cells than wild type or single knockout controls. We next examined the distinct cellular populations that were infected with Mtb in each genotype. We found that over 40% of infected cells in *Cybb*^*-/-*^*Caspase1/11*^*-/-*^ mice were found to be CD11b^+^GR1^int^ granulocytes a significant increase compared to wild type, *Cybb*^*-/-*^ and *Caspase1/11*^*-/-*^ animals (Figure 5E and 5F). This represents a shift in the *in vivo* intracellular distribution of Mtb in *Cybb*^*-/-*^*Caspase1/11*^*-/-*^ mice. Altogether these experiments show that the susceptibility of *Cybb*^*-/-*^*Caspase1/11*^*-/-*^ mice is associated with the recruitment of permissive granulocytes to the lungs that harbor high levels of Mtb.

## Discussion

While the phagocyte oxidase is undoubtedly important for protection against Mtb, the precise mechanisms by which it protects remain unclear (18, 21, 35, 36). In animal models, the loss of *Cybb* alone results in a loss of disease tolerance through increased Caspase1 activation (18). Our results show that phagocyte oxidase also contributes to protection through a mechanism that is revealed only in the absence of Caspase1/11. While loss of either *Cybb* or *Caspase1/11* results in minor changes in survival, combining the mutations resulted in a dramatic increase in susceptibility, similar to mice lacking *IFNγ, Nos2* or *Atg5* (7, 16, 17, 31). The synthetic susceptibility phenotype was characterized by increased granulocyte influx and Mtb replication in the lungs. Whether this susceptibility is a result of failed antimicrobial resistance, failed tolerance or both remains to be fully understood. However, based on the genetic interaction, it is likely that *Cybb* and *Caspase1/11* control parallel pathways that regulate cytokine and chemokine production and contribute to protection against TB.

While Mtb infection of both *Cybb*^*-/-*^ and *Cybb*^*-/-*^*Casp1/11*^*-/-*^ mice drives increased granulocyte trafficking to the lungs, the properties of these cells are distinct. In *Cybb*^*-/-*^ mice the granulocytes express high levels of GR1 and the distribution of Mtb infected cells is unchanged compared to wild type mice. In contrast, granulocytes recruited to the lungs of *Cybb*^*-/-*^*Caspase1/11*^*-/-*^ mice express intermediate levels of GR1 and are associated with high levels of Mtb. A recent report characterizing the susceptibility of mice deficient in *Nos2* found that GR1^int^ granulocytes were long-lived, unable to control bacterial growth, and were not suppressive even with increased IL10 production (33). In humans, low density granulocyte populations are associated with severe susceptibility to TB and may be analogous to these permissive GR1^int^ cells seen in susceptible mice (37). It is possible that these granulocytes are not directly driving the susceptibility but rather are associated with uncontrolled TB disease caused by other defects in the host response. Future work using depletion and conditional knockouts will be required to understand how these changes in the cellular dynamics in *Cybb*^*-/-*^*Caspase1/11*^*-/-*^ mice contribute to susceptibility.

Our current model predicts that the phagocyte oxidase and Caspase1/11 control the inflammatory state of myeloid cells during Mtb infection. When this control is lost, the result is a failure of disease tolerance, which drives progressive disease and recruits permissive granulocytes that modulate a feedforward loop of inflammation, Mtb growth, and tissue damage. While the exact signals that recruit granulocytes to the lungs of *Cybb*^*-/-*^*Caspase1/11*^*-/-*^ mice remain unclear, we observed dysregulation of IL6, CXCL2 and IL1α in Mtb infected *Cybb*^*-/-*^*Caspase1/11*^*-/*-^ macrophages. While the importance of each cytokine to *Cybb*^*-/-*^*Caspase1/11*^*-/-*^ susceptibility will need to be examined extensively, IL1α was the most significantly changed cytokine in *Cybb*^*-/-*^*Caspase1/11*^*-/-*^ macrophages and *in vivo*. IL1α is known to be required for protection against Mtb, as knockout mice are highly susceptible to disease (19, 38). Whether exacerbated IL1α directly contributes to TB susceptibility remains to be fully understood. Several non-mutually exclusive mechanisms could explain the dysregulation of IL1α and possibly other cytokines.

For example, there is evidence that changes in calcium influx and mitochondrial stability directly control the expression and processing of IL1α (39). Thus, Cybb or Caspase1/11 may modulate calcium flux and mitochondrial function during Mtb infection that activate excessive IL1α production. Recent studies also suggest that metabolic pathways control ROS production that is directly required for processing of GSDMD which may link cellular metabolism to ROS signaling and inflammasome function (40). There is also evidence of a direct interaction between the phagocyte oxidase subunits and Caspase1 that modulates phagosome dynamics during *Staphylococcus aureus* infection, but if this mechanism plays a role during Mtb infection remains unknown (41). Ongoing work is focused on examining the contribution of each potential mechanism to the susceptibility of *Cybb*^*-/-*^*Caspase1/11*^*-/-*^ animals to better understand the regulatory networks that control inflammatory TB disease.

One outstanding line of questions from our findings is the specificity of the genetic interaction between *Cybb* and *Caspase1/11*. Both Caspase1- and Caspase11-dependent pathways are activated during Mtb infection, yet the direct contribution of either Caspase1 or Caspase11 to our observed susceptibility remains to be investigated by individually generating either *Cybb*^*-/-*^*Caspase1*^*-/-*^ or *Cybb*^*-/-*^*Caspase11*^*-/-*^ animals (18, 42-44). Given the recent availability of clean *Caspase1* and *Caspase11* knockout mice, we are in the process of developing these models for the future (45-47). In addition, whether mutations in the inflammasome sensor NLRP3 or the adaptor ASC and other subunits of the phagocyte oxidase recapitulate the susceptibility of *Cybb*^*-/-*^*Caspase1/11*^*-/-*^ remain unknown. Further dissecting these specific genetic interactions between other phagocyte oxidase and inflammasome components will help to elucidate the underlying mechanisms controlling the susceptibility observed in *Cybb*^*-/-*^*Caspase1/11*^*-/-*^ animals.

Our discovery of a synthetic susceptibility to Mtb in *Cybb*^*-/-*^*Caspase1/11*^*-/-*^ mice was serendipitous. As the susceptibility observed in *Cybb*^*-/-*^ mice was found to be due to dysregulated Caspase1 activation and IL1β production, we initially hypothesized that the combined loss of *Cybb* and *Caspase1* would reverse the tolerance defects found in mice lacking the phagocyte oxidase and were surprised to uncover a synthetic susceptibility. Since the phagocyte oxidase and inflammasomes are among the most studied pathways in immunology, our findings highlight a fundamental lack of understanding of interactions between immune signaling networks that control inflammation and immunity. To develop new host-directed therapeutics that could shorten treatment times and improve disease control, it is critical to understand how these interconnected networks function to protect against TB. A major obstacle in identifying protective networks against *Mtb* is the redundancy among host pathways, which mask important functions in single-knockout animals. A global understanding of genetic interactions that impact key inflammatory networks during TB would significantly inform the development of effective host-directed therapies or immunization strategies. Large-scale genetic interaction studies are common in cancer biology and should be applied to immune signaling networks during Mtb infection to better define these critical but currently unknown mechanisms that control protection against TB (48). Altogether, these findings suggest genetic interactions are key regulators of protection against Mtb with Cybb and Caspase1/11 contributing together to protect against TB.

## Materials and methods

### Mice and Ethics Statement

Mouse studies were performed in accordance using the recommendations from the Guide for the Care and Use of Laboratory Animals of the National Institutes of Health and the Office of Laboratory Animal Welfare. Mouse studies were performed using protocols approved by the Institutional Animal Care and Use Committee (IACUC) in a manner designed to minimize pain and suffering in *Mtb*-infected animals. All mice were monitored and weighed regularly. Mice were euthanized following an evaluation of clinical signs with a score of 14 or higher. C57BL6/J mice (# 000664) and *Cybb*^*-/-*^ mice (# 002365) were purchased from Jackson labs. *Caspase1/11*^*-/-*^ were a kind gift from Katharine Fitzgerald and *Cybb*^*-/-*^ *Caspase1/11*^*-/-*^ were generated in-house. All mice were housed and bred under specific pathogen-free conditions and in accordance with the University of Massachusetts Medical School (Sassetti Lab A221-20-11) and Michigan State University (PROTO202200127) IACUC guidelines. All animals used for experiments were 6-12 weeks old.

### Macrophage and dendritic cell generation

Bone marrow-derived macrophages and dendritic cells were obtained from the femurs and tibias of sex- and age-matched mice. For BMDMs, cells were cultured in 10cm^2^ non-tissue culture treated petri dishes with 10 mls DMEM with 10% FBS and 20% L929 supernatant for 1 week. On day 3, the old media was decanted, and fresh differentiation media was added. After 7 days of differentiation, cells were lifted in PBS with 10mM EDTA and seeded in tissue-culture treated dishes in DMEM with 10% FBS with no antibiotics then used the following day for experiments.

For BMDCs, cells were cultured in 10cm^2^ non-tissue culture treated petri dishes with 10 ml DMEM with 10% FBS, L-Glutamine, 2 µM 2-mercaptoethanol and 10% supernatant from B16-GM-CSF cells as described previously. After 7 days of differentiation, BMDCs were further enriched by isolating loosely adherent cells removing F4/80^+^ cells then isolating CD11c^+^ cells by bead purification following manufacturer’s instructions (Stem Cell Tech). Cells were then plated in tissue culture treated dishes in DMEM with 10% FBS then used the following day for experiments.

### Bone marrow-derived macrophage and dendritic cell infections and analysis

PDIM positive H37Rv was grown in 7H9 medium containing 10% oleic albumin dextrose catalase growth supplement and 0.05% Tween 80 as done previously (18). Prior to infection, cultures were washed in a PBS-0.05% Tween solution and resuspended in DMEM with 10% FBS. To obtain a single cell suspension, samples were centrifuged at 200xg for 5 minutes to remove clumps. Culture density was determined by taking the supernatant from this centrifugation and determining the OD_600_, with the assumption that OD_600_ = 1.0 is equivalent to 3×10^8^ bacteria per ml. Bacteria were added to macrophages for 4 hours then cells were washed with PBS and fresh media was added. For cytokine analysis, at the indicated time points, supernatants were harvested and centrifuged through a 0.2-micron filter.

Supernatants were then analyzed by a Luminex multiplex assay (Eve Technology) or by ELISA following manufacturer protocols (R&D). For CFU analysis, at the indicated timepoints, 1% saponin was added to each well without removing media to lyse cells while maintaining extracellular bacteria. Serial dilutions were then completed in phosphate-buffered saline containing tween 80 (PBS-T) and dilutions were plated on 7H10 agar. For cell death experiments, at the indicated time points media was removed and a CellTiter-Glo assay (Promega) was completed following manufacturer’s instructions.

### Mouse infections and CFU quantification

For animal infections, H37Rv or YFP^+^ H37Rv were resuspended in PBS-T. Prior to infection, bacteria were sonicated for 30 seconds, then delivered into the respiratory tract using an aerosol generation device (Glas-Col). To verify low dose aerosol delivery, a subset of control mice was euthanized the following day. Otherwise the endpoints are designated in the figure legends. To determine total CFU in either the lung or spleen, mice were anesthetized via Carbon Dioxide asphyxiation and cervical dislocation. the organs were removed aseptically and homogenized. 10-fold serial dilutions of each organ homogenate were made in PBS-T and plated on 7H10 agar plates and incubated at 37C for 21-28 days. Viable bacteria were then counted. Both male and female mice were used throughout the study and no significant differences in phenotypes were observed between sexes.

### Flow Cytometry

Analysis of infected myeloid cells in the lungs was done as previously described (13, 33). In short, lung tissue was homogenized in DMEM containing FBS using C-tubes (Miltenyi). Collagenase type IV/DNaseI (Sigma) was added, and tissues were dissociated for 10 seconds on a GentleMACS system (Miltenyi). Lung tissue was then oscillated for 30 minutes at 37C. Following incubation, tissue was further dissociated for 30 seconds on a GentleMACS. Single cell suspensions were isolated following passage through a 40-micron filter. Cell suspensions were then washed in DMEM and aliquoted into 96 well plates for flow cytometry staining. Non-specific antibody binding was first blocked using Fc-Block. Cells were then stained with anti-GR1 Pacific Blue, anti-CD11b PE, anti-CD11c APC, anti-CD45.2 PercP Cy5.5 (Biolegend). Live cells were identified using zombie aqua (Biolegend). No antibodies were used in the FITC channel to allow quantification of YFP^+^ Mtb in the tissues. All experiments contained a non-fluorescent H37Rv infection control to identify infected cells. Cells were stained for 30 minutes at room temperature and fixed in 1% Paraformaldehyde for 60 minutes. All flow cytometry was run on a MACSQuant Analyzer 10 (Miltenyi) and was analyzed using FlowJo version 9 (Tree Star).

### Statistical Analysis

Statistical analyses were performed using Prism 10 (Graph Pad) software as done previously (18, 49). Statistical tests used for each experiment are described in each figure legend along with symbols indicating significance or no significance.

## Acknowledgements

We thank members of the Olive lab and Christopher Sassetti for help discussions. We thank the Fitzgerald lab for sharing *Caspase1/11*^-/-^ mice. This work was funded by National Institutes of Health grants AI148961 and AI165618 to Andrew Olive.

